# Phototaxis of the dominant marine pico-eukaryote *Micromonas sp*.: from population to single cell

**DOI:** 10.1101/740571

**Authors:** Richard Henshaw, Raphaël Jeanneret, Marco Polin

## Abstract

*Micromonas commoda* (previously *Micromonas pusilla*, a unicellular photosynthetic picoeukaryote globally dominant in marine ecosystems, has previously been qualified as being strongly phototactic. To date, no detailed quantitative or qualitative description of this behaviour has been reported, nor have thorough studies of its motility been undertaken. This primary producer has only been qualitatively described as utilizing run-and-tumble motion, but such motility strategy is incompatible with its morphology comprising only one propelling flagellum. Moreover, it is still unclear as to how *Micromonas sp.* detects a light direction due to the lack of a dedicated eyespot; the organism is essentially blind. Here we first perform population-scale phototactic experiments to show that this organism actively responds to a wide range of light wavelengths and intensities. These population responses are well accounted for within a simple drift-diffusion framework. Based on single-cell tracking experiments, we then detail thoroughly *Micromonas sp.*’s motility which resembles run-and-reverse styles of motion commonly observed in marine prokaryotes and that we name *stop-run or reverse*. The associated peculiar microscopic changes upon photo-stimulation are finally described and integrating those into jump-diffusion simulations appears to produce phototactic drifts that are quantitatively compatible with those obtained experimentally at the population level.

## INTRODUCTION

The fitness of aquatic microbes strongly depends on their ability to locate and aggregate at food sources known as being dynamically patchy on the micro-to the meso-scale [1–3] and at the same time to avoid hazardous situations (predators, parasites, etc) [4]. Consequently, most pelagic micro-organisms have accumulated extensive physico-bio-chemical toolboxes enabling them to move in the water column, either by using swimming apparatus such as flagella [5] or by controlling their buoyancy [6], and to sense adequately their environment and dynamically respond to it in order to find stimulating growth conditions. Such behaviors are mediated by multitude forms of *taxis*, whereby cells actively or passively reorient themselves and rectify their otherwise isotropic random motion in external scalar or vectorial fields such as chemicals (chemotaxis) [2, 3, 7], lights (phototaxis) [8–10], fluid velocities (rheotaxis) [11, 12], gravitational fields (gravitaxis) [13, 14], magnetic fields (magnetotaxis) [15, 16] or even fluid viscosities (viscotaxis) [17, 18].

Phytoplanktonic species are no exception. These prokaryotic and eukaryotic photosynthetic organisms constitute the basic food reservoir of higher trophic species that benefit from their high-conversion rates between solar energy and carbon-based molecules: they contribute to an estimated ∼ 50% of primary carbon production despite making up only ∼ 1% of the Earth biomass [19, 20]. As it happens, their very high photosynthetic yield explains the recent development of a plethora of algal factories for biofuel and food production [21]; such industry imposing also negligible pressure on arable land. Among the known ∼ 5000 species (only in marine ecosystems) [22], a few are thriving significantly more than the vast majority. This includes notably species from the genus diatoms, cyanobacteria, dinoflagellates and green algae. Understanding the reasons why those cope so well with their environment, how their evolution and adaptation have led to their success and how this picture can change in our climate-changing world appears therefore essential, from an ecological and economical perspective.

In this context, the green alga *Micromonas commoda* (previously *Micromonas pusilla* until a recent reclassification of the species in the *Micromonas genus* [23]) has a great role to play as it is one of the most abundant species in the ocean and is found in marine and coastal environments globally, from the Caribbean Sea to the Arctic waters [24–26]. Since the seminal work of Manton and Parke in 1960 [27] who described in detail its morphology and some qualitative features of its motility, important advances regarding the biology and ecology of this species have been established [7, 28–32]. Surprisingly though, our current knowledge regarding its motility, foraging strategy and single-cell response to stimuli is still very limited. From [27] we know that its ∼ 1 − 2*µ*m comma-shaped body (the smallest known eukaryote on Earth!) is complemented by a single ∼ 5*µ*m flagellum which protrudes from the concave side of the cell and propels it up to ∼ 50 body lengths per second [8]. *Micromonas sp.* apparently changes its swimming direction frequently [8] but the way it does so with a single flagellum remains unclear. More recently Omoto *et al* [33] have shown that contrary to most eukaryotic flagella that bend and propagate elastic waves, *Micromonas sp.* flagellum rotates similarly to bacteria. Moreover, given a particular orientation of the body, the cell can swim in two opposite directions [27], probably due to opposite rotations of the flagellum like the run-reverse-flick bacterium *Vibrio alginolyticus* [34]. Finally, *Micromonas sp.* has been qualitatively described as being strongly phototactic [27] which is common among photosynthetic microorganisms [8–10], but the range of light intensities and wavelengths that trigger this behavior as well as the single-cell dynamics leading to it are still unknown. This phototaxis is all the more surprising that this species does not have a dedicated eyespot to detect light, which suggest that such behavior is strongly coupled to the photosynthetic activity.

In this study we propose to start filling this gap by shedding light on the motility and single-cell phototaxis of this globally important species. We first use time lapse microscopy to show that *Micromonas sp.* indeed exhibits net positive phototaxis at the population level under a wide range of light wavelengths and intensities. We then perform single-cell tracking experiments to thoroughly describe the foraging strategy of non-stimulated cells and to understand how these microscopic features are altered upon photo-stimulation. Finally, in order to relate our findings from single-cell dynamics to population phototaxis, we carry out simulations that integrate the microscopic dynamics of the cells and are seen to produce phototactic drifts in quantitative agreement with our population experiments.

## METHODS

We briefly summarise here our experimental and numerical methods, and refer the reader to the Supplementary Information (SI) for a detailed description of our procedures.

### Culturing

Initial cultures of *Micromonas sp.*strain RC827 were generously provided by Dr Joseph Christie-Oleza. These cultures were grown and swapped between two medium to promote cell growth - Keller (K) medium [35] and Gulliard’s f/2 medium [36], prepared using artificial seawater in 500 ml quantities excluding the sodium glycerophosphate to reduce precipitation. The cultures were cultivated in a diurnal chamber on a 16*/*8 hour light/dark cycle, maintained at 20°C at a light intensity of 100*µ*E.m^−2^.s^−1^ from fluorescent strips. All experiments took place with the cells growing in f/2 medium.

### Experimental Setup

A sketch of the experimental setup is shown in Fig. 3a. All experiments were done using a Nikon TE 2000U inverted microscope fitted with a longband pass filter (765nm cutoff wavelength, Knight Optical UK) in order to limit phototactic stimulation from the imaging light, a 40× air objective and filmed using an Allied Vision Pike F-100B camera. Camera settings were optimized to reduce the required intensity of the optical beam to limit thermal convection. A cuboid polydimethylsiloxane (PDMS) chamber was cast with internal dimensions 10 mm ×10 mm×4 mm with a wedge insert added on the far edge of the chamber to reduce any coherent back-reflection, then bonded to a glass cover slip using a Harrick oxygen plasma cleaner. The experimental window covered 185 *µ*m from the boundary of the chamber closest to the stimulus and along the full width of the chamber. Prior to filming, the sample had been left to sediment for a period of several hours. Light stimulus was generated from one of four LEDs (then collimated) from Thorlabs with peak wavelengths *λ* = 470 nm, 530 nm, 595 nm and 625 nm. Light intensity was measured with a Spectrum Technologies Field Scout light sensor reader.

**FIG. 1.**
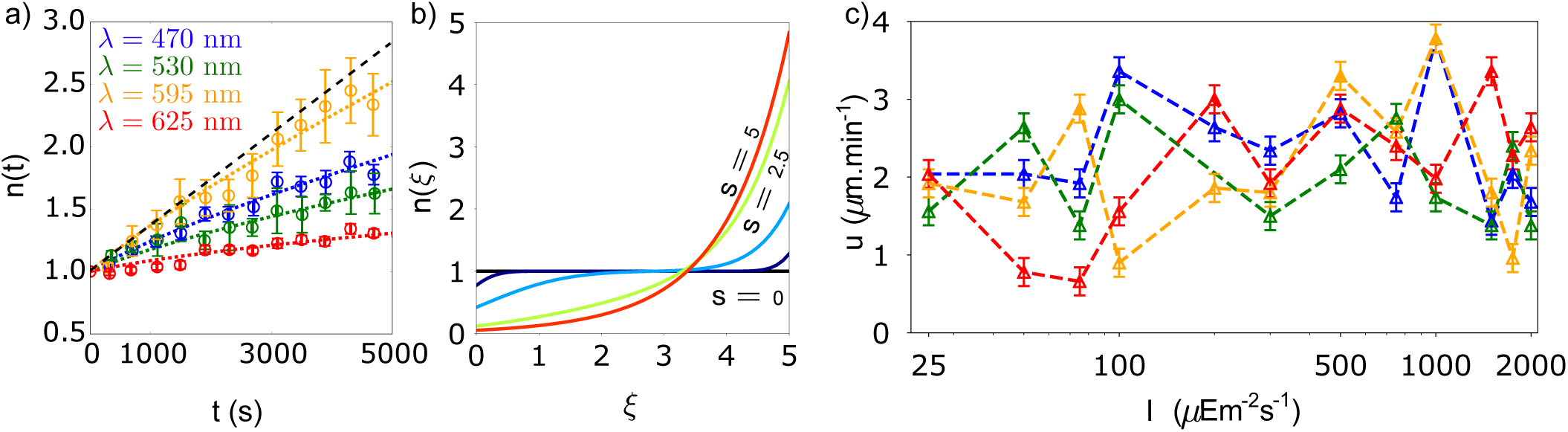
Population-scale phototaxis of *Micromonas sp.*. **a)** Evolution of the number of cells (normalized by the initial number) within the field of view close to the light source at *I* = 75*µ*E.m^−2^.s^−1^ for the four wavelengths examined (circles) together with the best fit from the drift-diffusion solution (dotted lines). Black dashed line is a linear extrapolation of the initial population growth where drift dominates diffusion. **b)** Time-evolution of the drift-diffusion model (Eq 1) starting from a uniform distribution of cells across the 1D system (black curve). The solution converges towards an exponential steady-state (red curve). We show the distribution *n*(*ξ*) for 5 dimensionless times *s* = 0, 0.05, 0.5, 2.5 and 5 (*ξ* is a dimensionless spatial coordinate, see SI §1 for a detailed calculation). **c)** Phototactic response quantified by the drift velocity *u* as a function of intensity *I* for the four wavelengths examined. The phototactic drift *u* is always positive showing a positive phototactic response for every parameter tested.

**FIG. 2.**
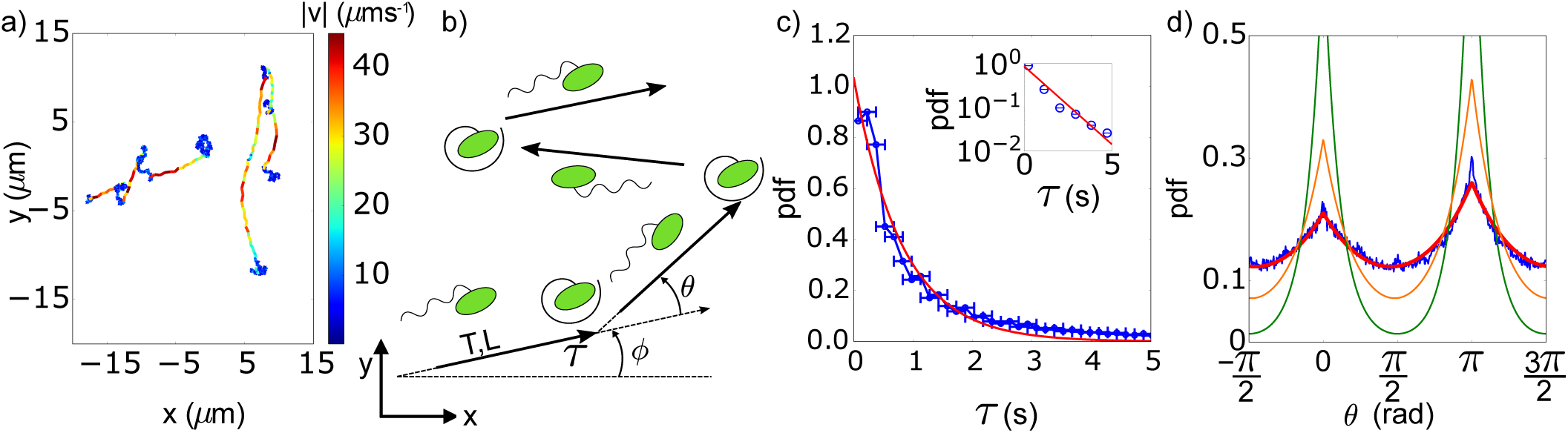
Micromonas sp. motility. **a)** Examples of *Micromonas sp.* trajectories, colour-coded by the instantaneous velocity (see colormap). The cells alternate between stop phases (blue-ish parts) and forward or backward runs (red-ish parts). **b)** Stop-run or reverse schematic. The cell runs for a length *L* (during a time *T*) at an angle *ϕ* relative to the *x*-axis, reorients during a time *τ* after which the cell moves again in a direction *ϕ* +*θ*. Displayed changes in the flagellum configuration are purely to aid visualisation as the physical flagellum behaviour is unknown. **c)** Distribution of waiting times *τ* relatively well fitted by a simple exponential. Inset: semi-log plot of the distribution. **d).** Distribution of reorientation angle *θ*. The distribution is bimodal with peaks at *θ*_run_ = 0 and *θ*_rev_ = *π*; the cells either continue in the current direction or a reversal takes place after a stop. The experimental data (blue) is perfectly fitted with Eq. 3 (red, see also SI §3). Shown in orange (resp. green) is the theoretical distribution (Eq. 3) for a completely passive process with particle radius 0.63 *µ*m (resp. 1 *µ*m) (*D*_rot_ = 0.64 and 0.15rad^2^.s^−1^ respectively). The former is chosen as the equivalent spherical radius of the cell, see text.

**FIG. 3.**
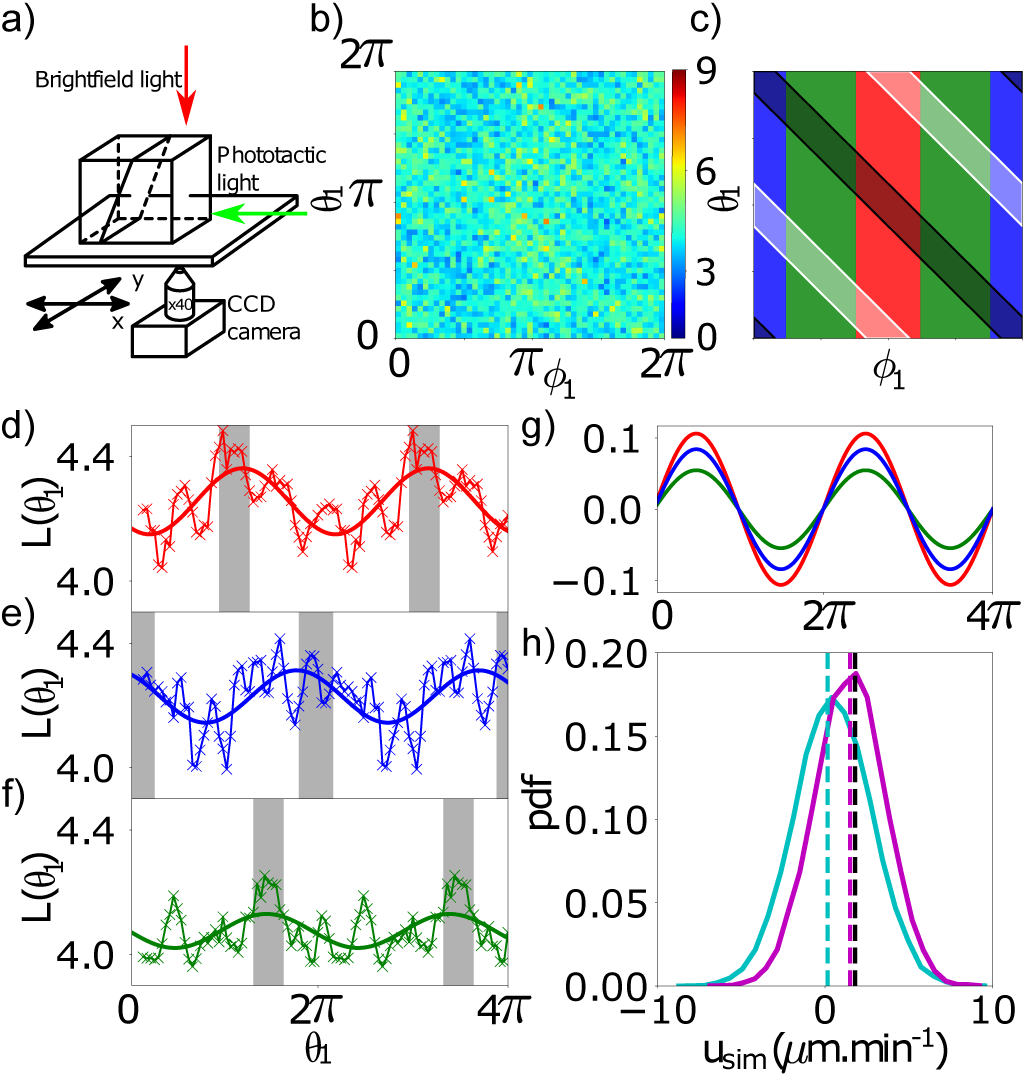
Individual cell phototaxis. **a).** Sketch of the experimental setup and incoming light direction denoting the *ϕ* = 0 run angle. **b).** Average run length as a function of previous run direction *ϕ*_1_ and reorientation *θ*_1_. **c).** Areas of interest in panel b). Previous runs away from (resp. towards/orthogonal to) the light source in red (resp. blue/green). Reorientation angles that bring the cells towards (resp. away from) the light are shown in black (resp. white). **d-f).** Run length *L*(*θ*_1_) averaged over *ϕ*_1_ in each of the three zones in panel c). Full lines are simple sinusoidal fittings to highlight the trend. The run length modulation is always maximum for angles *θ*_1_ bringing the cells towards the light (vertical grey shading). For a better visualisation of the modulation, we have concatenated the same 2*π*-signals over 4*π*. **g).** Sinusoidal fittings in panels c-e) after eliminating the phase and average values to show the difference in the modulation depending on the previous run direction *ϕ*_1_. The response is maximal (resp. minimal) when the cell first swims away from (resp. orthogonal to) the light, red curve (resp. green). **h).** Distribution of drift velocities (full lines) and average drift values (vertical dashed lines) in the jump-diffusion simulations with/without light (magenta/cyan respectively). The simulations with light reproduce very well the population drift for this set of intensity/wavelength (black dashed line).

### Population Experiments

20 equally spaced sites were sampled along the edge of the chamber nearest the light stimulus. Using a computer controlled microscope stage, the population for each site was counted in a cyclical manner for a duration of 88 minutes per site. These sites were then combined to give a total population profile versus time for the total experimental area with a sampling period of 230 s for each full cycle.

### Cell Motility Experiments

5, 000 frames films were taken with a framerate between 25 − 30 FPS at locations randomly chosen over the surface of the chamber. These frames were then pre-tracked and the particle trajectories reconstructed using MATLAB^®^ functions developed by Blair and Dufresne [37]. From these trajectories the individual cell motion parameters were measured. These experiments were repeated for both control cases and for stimulated cases where a 625 nm stimulus at 300 *µ*E.m^−2^.s^−1^ intensity was used. Two different stimulus exposure times were used: a shorter time scale of 0−45 minutes and 90−120 minutes of stimulation respectively. Films were taken at a height of 20 *µ*m above the coverslip to prevent cells from tethering to the surface with the flagellum and to reduce hydrodynamic interactions [38] so the cells swim approximately straight.

### Particle trajectories

A total of 480, 939 control tracks were analysed, with 385, 625 tracks consisting of a (minimum of) a complete run-reorient-run cycle referred to as tracks and reorientation tracks respectively. Two stimulated data sets were collected, firstly over a period of 45 minutes of constant stimulation (235, 152 tracks with 193, 146 reorientation tracks) and a second set during 90 − 120 minutes of constant stimulation (447, 526 tracks with 359, 967 reorientation tracks), totalling at 1, 163, 617 tracks analysed. Only tracks that could be approximated to linear were considered, and all trajectories required a minimum of 100 data points before analysis (i.e. corresponding to ∼ 3 − 4s of acquisition time).

### Jump-diffusion simulations

For both the control and phototactic cases 2, 000 particles were simulated for a duration *t*_end_ = 5400s with a timestep of Δ*t* = 0.004s. These numerical experiments were based on the 2D jump-diffusion simulations performed by Jeanneret *et al.* [39]. Particles were given a previous run direction *ϕ*_0_ and reorientation *θ*_0_ sampled randomly from [0, 2*π*[, which gives the initial run angle *ϕ*_1_. The jump length is sampled directly from the distribution of run lengths as a function of previous run angle/reorientation angle, as shown in Figure 3b. After each run, the particle samples a waiting time from the exponential distribution shown in Figure 2c during which it diffuses both rotationally (*D*_rot_ = 2.1rad^2^.s^−1^, see section Cell motility) and translationally (*D*_Brown_ = 0.34*µ*m^2^.s^−1^ taking an equivalent spherical radius *R* = 0.63*µ*m for the particle). The rotational diffusion over the waiting time together with the probability of reversal (given by Γ = 0.55 in Equation 3) then determine the reorientation angle *θ*_1_. From *ϕ*_1_, *θ*_1_, the run length for the next run can be sampled and the process repeats for the duration of the simulation. Diffusion is completely turned off during a run. Finally, we define the drift velocity for each simulated cell as *u*_sim_ = (*x*(*t*_end_) − *x*(0))*/t*_end_.

## RESULTS

### A. Population Phototaxis

To quantify the phototactic response of the population we first measure the accumulation of cells at the edge of our chamber nearest to the light source (Fig. 3a) over 88 minutes exposure time for a large set of light intensities and wavelengths. The evolution of cell number in that region, *n*(*t*) = *N* (*t*)*/N*_0_ with *N*_0_ the initial number, is plotted Fig. 1a for the four wavelengths examined and at fixed intensity *I* = 75*µ*E.m^−2^.s^−1^. Clearly, an accumulation of cells is observed for each wavelength that was probed, showing a positive phototactic response at this particular intensity. Because *Micromonas sp.* displays diffusive motion on long-time scales due to stochastic re-orientations (see next sections), we expect the accumulation dynamics to be well described by a 1D drift-diffusion model:

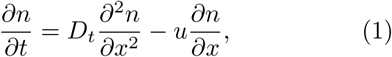

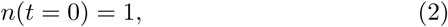

where the light comes from the positive x-direction, *u* is a population drift velocity modelling the phototactic response of the population (*u* > 0 for positive phototaxis) and *D*_*t*_ represents the translational diffusion coefficient of the cells. Across the width of the system the steady-state solution is exponential with a maximum at the boundary closest to the light source when *u* > 0 (red curve in Fig. 1b, see SI §1 for a detailed calculation of the theoretical solution). Fitting now the experimental data with this model (considering the theoretical solution of ⟨*n*(*t, x*) _*x*∈f.o.v._, see SI §1)⟩ gives a remarkable agreement (Fig. 1a) when leaving *u* as the sole free-fitting parameter and using *D*_*t*_ = 1.17 *µ*m^2^.s^−1^ (measured from single cell tracks, see next sections). The phototactic velocity *u* extracted from this procedure is shown to be positive for every wavelengths and intensities tested (Fig. 1c), synonymous with a phototactic response directed towards the light. The dependency of the phototactic response with light intensity and wavelength seems highly non-trivial and does not exhibit any particular trend. The only noticeable trait is the dimmer response to red light stimulation at low intensity compared to blue/green wavelengths (Fig. 1c).

Altogether we confirm quantitatively that *Micromonas sp.* is a strongly phototactic organism as previously noted qualitatively [27]. However, the magnitude of the response remains always quite low: the drift velocity never exceeds *u* ≈ 4 *µ*m.min^−1^. The object of the next sections is then to establish the microscopic origins of this phototactic response based on the characteristics of single cell motion with and without light stimulations.

### B. Cell motility: stop-run or reverse

We first characterise the motility of *Micromonas sp.* without light stimulation. Typical recorded tracks are presented Fig. 2a showing that the microalga alternates between phases of active swimming in straight runs (redish parts) and phases of apparent passivity where reorientation takes place (blue-ish parts). At first sight such motility pattern looks akin to the run-and-stop motion of *Rhodobacter sphaeroides* [40, 41], but further quantitative analysis are needed to assess the similitudes and differences between the motility strategy of the two species. To that purpose we extract different parameters from the experimental tracks (see the methods of extraction in SI §2 and the schematic in Fig. 2b): the run time *T*, the run length *L*, the run direction *ϕ* given by the angle between the x-axis and the swimming direction, the stopping time *τ* and finally the reorientation angle *θ* between two consecutive runs.

The stochastic switching events from running to stopping states and inversely both follow Poisson processes as commonly found in run-and-tumble type of motion [42]. This is illustrated by the distributions of run times *T* (Fig. S3c) and stopping times *τ* (Fig. 2c and Fig. S3d) which are well fitted by simple exponentials with switching rates *κ*_run_ = 1*/*⟨*T*⟩ = 5 ± 1s^−1^ (i.e. ⟨*T*⟩ = 0.20 ± 0.04s) and *κ*_stop_ = 1*/* ⟨*τ*⟩ = 0.95 ±0.04s^−1^ (i.e. ⟨*τ*⟩ = 1.05 ± 0.04s). During a run, the organisms don’t swim at constant velocity (see color code in Fig. 2a)) but exhibit phases of acceleration and deceleration at the beginning and at the end respectively. This feature is probably associated to a conformal change of the flagellum at the onset and at the end of the propulsion as observed with *R, sphaeroides* [43]. Peak run speeds can be up to ∼ 80*µ*m.s^−1^ (Fig. S3a) while the average run speed is 22.89 ± 0.02*µ*m.s^−1^ (Table S1 for numerical values of different average motility parameters). This leads to an average run length of a few body lengths ⟨*L*⟩ = 4.54 ±0.01*µ*m (full distribution of *L* in Fig. S3b). Because *Micromonas sp.* is monoflagellated (like the bacterium *R. sphaeroides*) and therefore cannot tumble as multiflagellated organisms, we are primarily interested in quantifying the reorientation of the alga during a stop phase. As can be guessed from the tracks in Fig. 2a, the direction of a run after a stop is not random, but strongly correlated with the previous one: the distribution of re-orientation angle *θ* is bimodal with peaks at *θ*_run_ = 0 and *θ*_rev_ = *π* (Fig. 2c). The cells either keep swimming in the same direction or completely reverse their motion after a stop. This contrasts with *R. sphaeroides* which exhibits a distribution of reorientation angles simply peaked at *θ*_run_ = 0 [40]. Such motility behaviour constitutes a strong proof that *Micromonas sp.* must possess the morphological ability to swim in two opposite directions from a given orientation of its body (as already noted in [27]), although the mechanism by which it does so remains elusive.

As illustrated by the red solid line in Fig. 2d, the full shape of the distribution is very well described by the analytical expression given in Eq. 3 below. Such expression describes the reorientation process simply as the consequence of rotational diffusion over exponentially distributed stopping times *τ* (see SI §3 for further details):

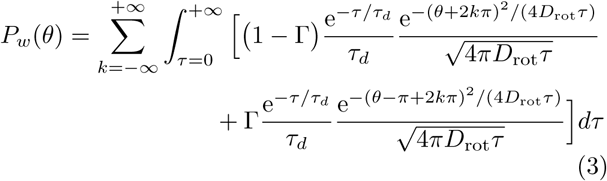

with Γ the probability to reverse from the previous run direction, *τ*_*d*_ the mean stopping time, and *D*_rot_ the rotational diffusivity of the cell. From the fitting we obtain Γ = 0.55 ± 0.01, which means that the cell has a larger probability to reverse its motion, *τ*_*d*_ = 0.80±0.01s in close agreement with the direct measurement of the average stopping times ⟨*τ*⟩ = 1.05 ± 0.04s, and *D*_rot_ = 2.1 ± 0.1rad^2^.s^−1^.

Such value of *D*_rot_ seems incompatible with being solely set by thermal forces. Considering an equivalent spherical radius of the cells *R*_geo_ = 0.63*µ*m (average semi-principal axes *a* = 1*µ*m and *b* = 0.5*µ*m) gives *D*_rot_ = 0.64rad^2^.s^−1^, leading to a theoretical distribution far-off the experimental one (using Eq. 3 and keeping *τ*_*d*_ = 0.80s and Γ = 0.55, orange curve in Fig. 2d). This discrepancy is expected to be even larger when taking into account the correction to the Brownian rotational diffusivity due to the non-spherical shape of the body (i.e through Perrin friction factors, see [41, 44]). Consequently, we conjecture that the reorientation process is sustained by some yet unknown active mechanism that enhances the rotational diffusivity during the stops. Based on a similar approach, the same conclusion was drawn from the microscopic features of the run-and-stop motility of *R. sphaeroides* [41]. The authors there hypothesized that this active contribution could originate from low-frequency motion of the flagellum during the stops [43] or from rapid polymorphic transformations of the flagellum during the stop-to-run transitions inducing rotation of the body [45]. We believe a similar process happens in *Micromonas sp.* although dedicated experiments aiming to record both flagellum and body motion would be needed in order to unequivocally reveal the re-orientation dynamics.

Overall this way of exploring the space leads to a translational diffusivity *D*_*t*_ ≈ 1.2*µ*m^2^.s^−1^ (obtained from the MSD, see SI §4), a few times the Brownian diffusivity of a sphere of similar size as *Micromonas sp.*. To the best of our knowledge, the motility pattern described here has no equivalent in previous literature. Because it appears somehow as a combination between the run-stop motion of the chemotactic species *R. sphaeroides* [40, 41, 46–48] and the run-reverse motility of *P. haloplanktis* [49–51], we coin this new foraging strategy *stop-run or reverse.*

### C. Individual Cell Phototaxis

To assess the origin of the phototactic drift seen at the population level, we repeat the single cell tracking experiment whilst imposing a continuous light stimulus from the positive *x*-direction (Fig 3a) at *λ* = 625nm and *I* = 300*µ*E.m^−2^.s^−1^. Data was collected between 0 and 45 minutes after switching on the light as well as between 90 and 120 to appraise any kinds of acclimation [52]. Surprisingly, none of the microscopic quantities introduced previously change upon light stimulation. In particular the distribution of reorientation angle *θ* remains identical to the control case (Fig. S5a), while the distribution of run angle *ϕ* remains uniform (Fig. S5b) showing no directed motion towards the light. Moreover we did not find any change in the run length distribution as a function of the run angle *ϕ*. The population drift is then the consequence of subtle spatial asymmetries in the statistical properties of the trajectories.

In fact, understanding the microscopic origin of the phototactic drift requires a thorough analysis of the complete run-stop-run pattern. We consider the average run length *L*(*ϕ*_1_, *θ*_1_) as a function of the previous run angle *ϕ*_1_ and reorientation angle *θ*_1_. The map of *L*(*ϕ*_1_, *θ*_1_) appears noisy and doesn’t seem at first sight to provide any clue regarding the phototactic behaviour of the cells (Fig. 3b, short phototactic stimulation experiment). To extract useful information, we first divide this map into three regions defined by the three colors in Fig. 3c where red (resp. blue, green) corresponds to the first run being oriented away from (resp. towards, orthogonal to) the light. The corresponding reorientation angles *θ*_1_ that bring the cell towards (resp. away from) the light are shown in shaded black (resp. white). Averaging over *ϕ*_1_ in each of the colored regions (e.g. 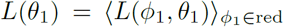) shows that the run length *L*(*θ*_1_) is modulated with respect to the reorientation angle *θ*_1_ (Fig. 3d-f, same color code), with a maximum (resp. minimum) when the cells run towards (resp. away from) the light in the second run (light direction=grey shading in Fig. 3d-f). To compare the modulation as a function of initial run direction *ϕ*_1_ we fit these curves with a simple sinusoidal function *L* = *γ* + *β* sin(*θ*_1_ − *δ*) (Fig. 3d-f, full lines). After eliminating the phase *δ* and subtracting the average value *γ*, the amplitude of the modulation is seen to be the largest when the cells swim away from the light in the first run, while it is the smallest when the cells swim first orthogonally to the light direction (Fig. 3g). In the control experiment, without light stimulation, no modulation of the run length is observed and the modulation response over long-time exposure is dimmed.

Finally, integrating our single-cell measurements with and without light into jump-diffusion simulations (see Methods) appears to produce drifts in good agreement with our population measurements. Indeed we obtain a distribution of drift velocity with average value ⟨*u*_sim_⟩ = 1.50 ± 0.02*µ*m.min^−1^ upon photo-stimulation, close to the value *u*_exp_ = 1.8 ± 0.3*µ*m.min^−1^ measured in this particular set of wavelength and intensity (Fig. 3h).

## DISCUSSION

While previous studies [27, 53, 54] have reported the existence of phototactic activity amongst the dominant marine pico-eukaryote *Micromonas sp.*, no clear picture of both the magnitude and the microscopic origins of this response has yet been established. Here by combining population-level and single-cell tracking experiments as well as numerical simulations, we have shown that this globally important species [24–26, 29, 55–57] is capable of regulating its motility in response to directed light stimuli, thus confirming the presence of phototaxis.

By examining a total of 1,163,617 single-cell tracks with and without light stimulation, we have shed lights on both the strategy *Micromonas sp.* adopts to explore its environment and the way it manages to slowly drift towards a light source. Interestingly, the cells perform a type of motion never reported before and that we dubbed *stop-run or reverse* (although a similar motility pattern has been reported in *E. coli* following forced cell body elongation [58]). This name reflects the fact that the cells frequently stop in order to partially reorientate via seemingly enhanced-rotational diffusion while they “choose” afterwards to move forward (‘run’) or backward (‘reverse’) with respect to their body orientation. Despite variants of run-reverse types of motion being common amongst marine prokaryotes [34, 49, 59], this is to the best of our knowledge the first reported example amongst eukaryotic species.

This strategy is certainly the consequence of the rudimental propulsive machinery of the organism (i.e. a single rotating flagellum [33]) which does not allow the cell to actively steer towards any controlled direction (nonetheless our data seems to show that the stochastic reorientation process is enhanced by some type of cell activity) as opposed to most algae that are also aided by dedicated eyespots for detecting light (e.g. *C. reinhardtii* [60], *Euglena gracilis* [61]) Instead, the phototactic drifts experienced by *Micromonas sp.* population originates from subtle directional asymmetries in run lengths. After reorienting back towards (resp. away from) the light source, the cell always

By examining the average run length for given values of the reorientations as a function of the incoming run angle we observe an increase (resp. decrease) in the runs that are reorientated back towards (resp. away from) the light source. This peculiar strategy is validated by encoding the experimental motility data into jump-diffusion simulations and measuring the simulated population drift this produces.

These observations probably reflect the way that *Micromonas sp.* manages to detect where the light comes from without a dedicated eyespot. Although the following mechanism is still hypothetical, the phototactic response could plausibly be mediated by the photosynthetic activity of the cell which takes place in the chloroplast located in a plastid in the nose of the organism [27]. In such case, the decrease (resp. increase) of photosynthetic activity during a run away from (resp. towards) the light might trigger an increase (resp. decrease) of the swimming speed as the organism reverses. In the case where the cell runs in a path quasi-perpendicular to the stimulus direction the photosynthetic activity would not change substantially, preventing the organism to effectively determine the direction of the light stimulus. This could explain why in the case where the previous run is perpendicular to the light source, the increase in the run length is reduced in comparison to runs that were initially on the light axis. This is supported by the observation of micro-lensing within the cell (SI 7) where we see the cell is capable of internally focussing light, which when coupled with a photorecpetor or measurements of the photosynthetic activity provides a viable phototactic steering mechanism reminiscent of cyanobacteria [62]. However despite being a plausible scenario given our experimental data, further investigation should aim to understand the complex biochemical processes taking place upon photostimulation as well as the flagellar apparatus dynamics.

The phototactic behaviour reported here only leads to relatively faint drifts with a maximum recorded speed at the population level *u* ≈ 4*µ*m.min^−1^. This contrasts with previous studies on chemotactic responses of this species to DMSP and derivatives where drifts were seen to be of the order of ∼ *µ*m.s^−1^ [7]. We believe this phototactic strategy is in fact mainly a way to counter-act sedimentation in order to remain at the same position in the water-column. Indeed the magnitude of the phototactic drifts obtained in this study matches well our measurement of the sinking speed of the cells when no light stimulus is imposed anywhere (SI). Importantly, the phototactic response is always directed towards the light source regardless of the wavelength or intensity, even when it could have been expected to be harmful for the cell (for instance via photo-oxidative stress [63, 64]). This contrasts with other phototactic organisms such as *C. reinhardtii* which switch to photophobic behaviours at high intensities in order to prevent self-damages [9, 65]. Although the response is always positive, the phototactic drift *u* does not follow any scaling law with the wavelength (i.e the curves in Fig. 1b are not parallel), which suggests that the cells make use of several distinct phytochrome photoreceptors [32, 66] whose activations depend on light intensity. Finally, we observed that *Micromonas sp.* is less sensitive to red than blue/green wavelengths at low intensities (*<* 100 *µ*E.m^−2^.s^−1^). This result probably reflects the natural condition of light in the oceans, where red wavelengths do not penetrate to depths greater than a few meters [67]. Consequently the peak sensitivity at low intensity would be shifted towards smaller wavelengths in order for the cells to keep track of the light when deep enough in the water layer.

## ACKNOWLEDGEMENTS

We would like to thank Dr Joseph Christie-Oleza for kindly supplying the initial cultures and Mr Steven Hindmarsh for technical assistance. RH acknowledges EPSRC Award 1619257 for funding.

## AUTHOR CONTRIBUTIONS STATEMENT

RH and MP designed the study; RH performed the experiments and data analysis; RJ and RH performed the simulations; RH, RJ and MP analysed the results and wrote the manuscript.

## ADDITIONAL INFORMATION

The author(s) declare no conflicting interests.

## Supplementary Material

### SI 1: SI DRIFT-DIFFUSION POPULATION BEHAVIOUR

Here we outline the main results from [S68], as discussed briefly in the main text. This is included for completeness and for accessibility to the material to the reader, for full methodology and mathematical support please see the source material.

Consider a colloidal suspension whose number density at a time *t* is given by *n*(*x, t*) with the *x*-coordinate restricted to the interval 0 ≤*x* ≤*h*. Given constant drift velocity *u* and Einstein coefficient *D*, the particle current *J* is described by

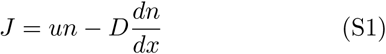

The density and current satisfy the conservation equation and also the zero current condition at the boundaries

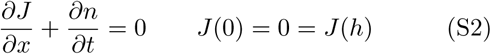

Combining these, we obtain the diffusion equation for the number density

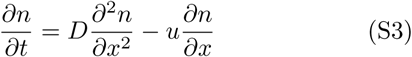

For convenience, we introduce the dimensionless variables

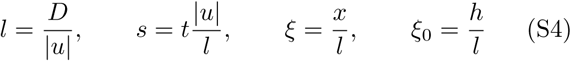

where *ξ*_0_ is in fact the Péclet number of particle diffusion. Using these variables, our original diffusion equation becomes

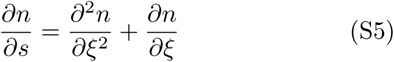

with *u <* 0 so the particles are driven towards the left boundary at *x* = 0. This direction of current is chosen arbitrarily, we could equally choose *u* > 0 such that the current is driven to the right boundary of *x* = *h*, in which case we simply set 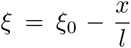 and continue with the methodology.

For the general initial distribution of particles, define the Green’s function of the process *g*(*ξ, ξ*_1_; *s*) which is the probability for a particle located position *ξ*_1_ at time *s* = 0 to be found at position *ξ* at time *s*. From this, the time evolution of the initial distribution *n*(*ξ*, 0) is described by evaluating

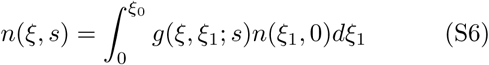

Substituting Equation S6 into Equation S5, the Green’s function can be shown to satisfy the dimensionless diffusion equation (Equation S5).

Solving for the Green’s function is a fairly arduous process, but the final form is given by

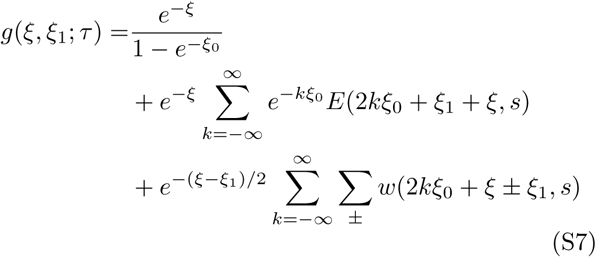

where the functions *E, w* are defined as

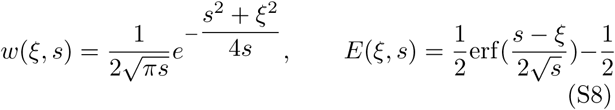

In the long time limit, the Green’s function tends to the stationary probability distribution

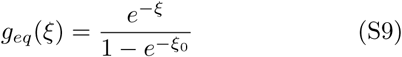

**FIG. S1.**
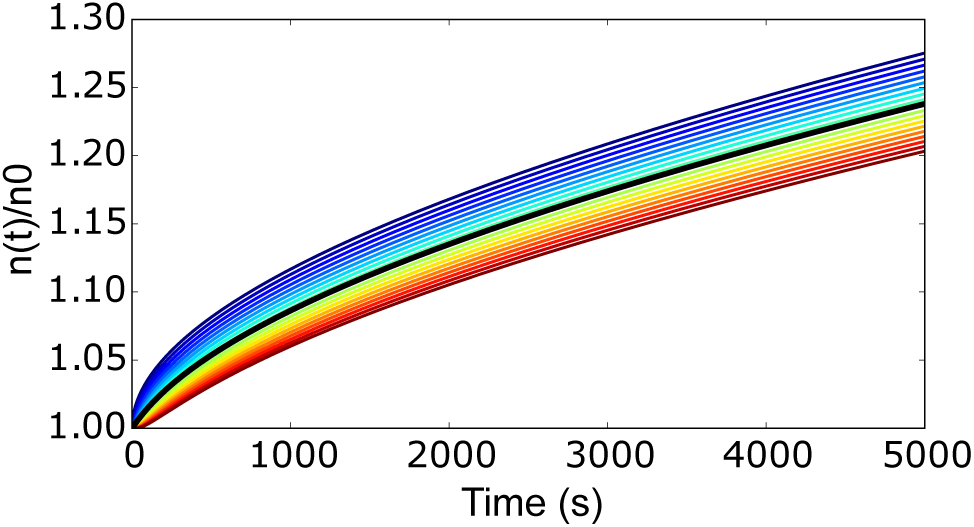
Drift-Diffusion model near the boundary. Population accumulation relative to an initial population *n*0 across the length of the experimental window. We take the average value of the population within 185 *µ*m of the boundary (our experimental window), labelled as the black line, to fit the population drifts in the main text. As expected the accumulation decreases as distance from the boundary increases.

For an initial constant density *n*(*ξ*, 0) = *n*_0_, Equation S6 becomes

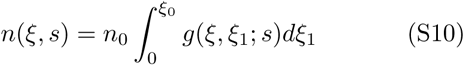

By integrating the Green’s function term by term, an expression for the time evolution of a flat original density can be found as

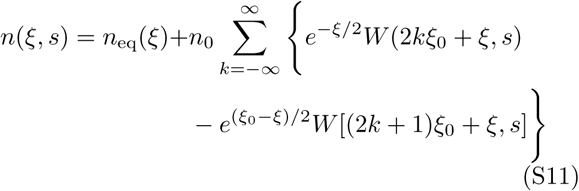

Where the function *W* and equilibrium value *n*_eq_ are defined as

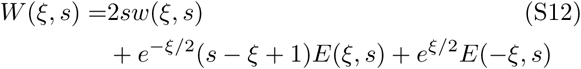

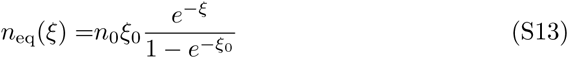

Finally, changing back to the original variables brings the equilibrium distribution to the form

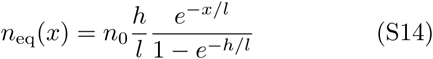

**FIG. S2.**
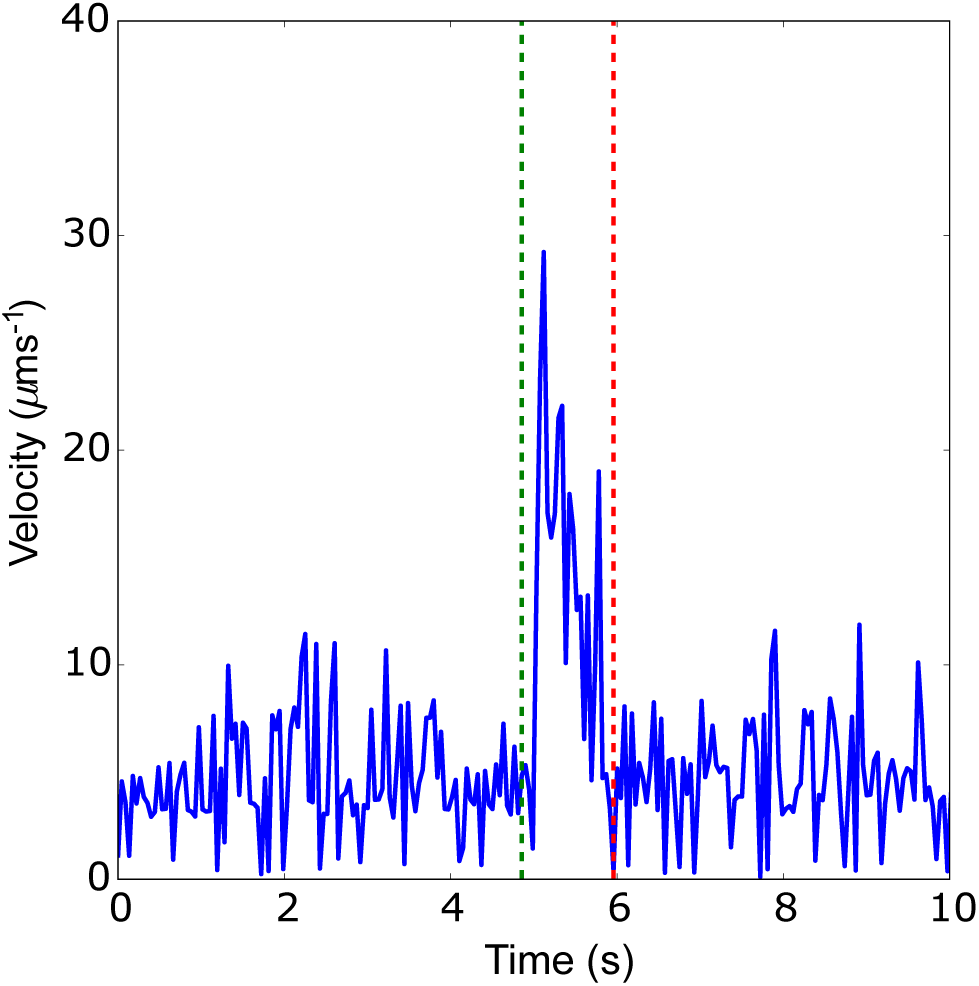
Run detection from instantaneous velocity. Run-events are easily identified from the background Brownian noise as sustained periods of time at higher speed. From identifying the initial peak, the start and end of the run can be determined (plotted as green and red respectively). From these start and end points the full run trajectory can be analysed to determine features such as the run length, run duration and run angle.

An example of the resultant population behaviour is shown in Fig. 1b-main text, where the population density converges to the exponential equilibrium distribution *n*_eq_. Figure S1 shows the evolution of the number density *n*(*t*) at fixed distances *x* ∈ [0, 185]*µ*m (blue to red, here 185*µ*m corresponds to the width of our field of view). As the distance from the boundary increases there is a decrease in the population accumulation at that point. This behaviour is expected as the particles at these points still have a distance they can travel before reaching the boundary. To take this behaviour into account when fitting the drift velocities we fit the experimental data to *N*_*B*_(*t*) defined as:

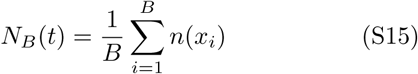

where *N*_*B*_ is evolution of the average number density of the population within a fixed distance of the boundary, where *B* is determined by the size of the observation window (185*µm* in our case). This is demonstrated in Figure S1 as the solid black line, denoting the average population value of the plotted range of *x* values.

### SI 2: CELL MOTILITY PARAMETERS

For a run-tumble organism it is possible to characterise the swimming behaviour with a selection of motility parameters: run length (L), run duration (T), tumble duration (*τ*), peak and average swimming speeds and the reorientation distribution (*θ*). The *θ* distribution has already been discussed in the main text (and below in section §3), so here we focus on the other motility parameters. The distributions of these are summarised in Figure S3 and the average values can be found in Table S1.

**FIG. S3.**
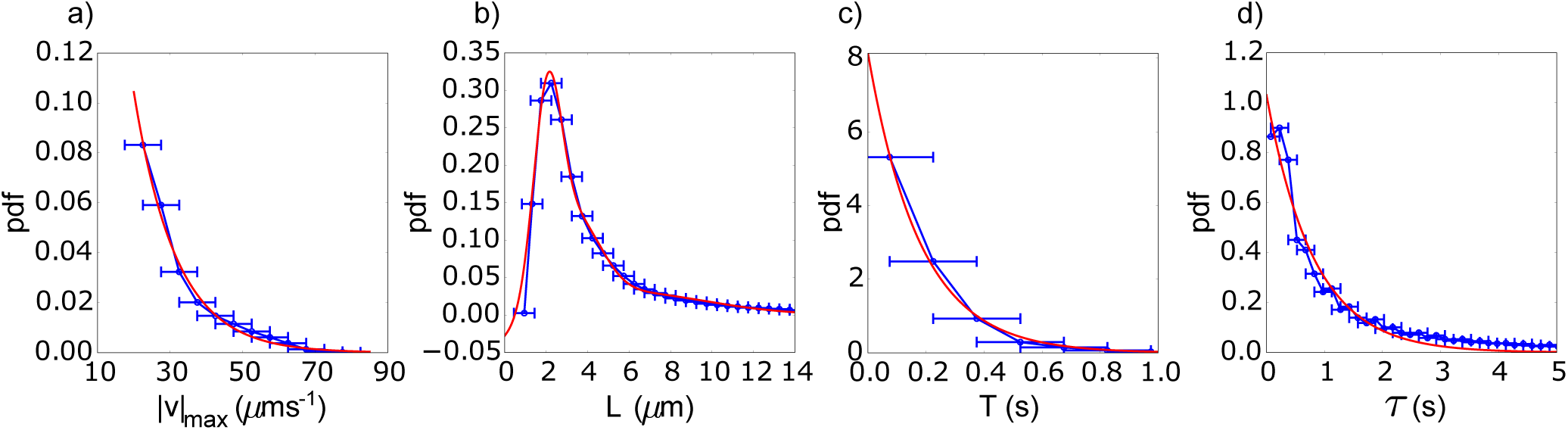
Observed cell motility parameters for *Micromonas sp.*. **a).** Peak run speed of individual trajectories. We have also observed extremely rare cases of the speed exceeding 100 *µ*ms^−1^ which are not recorded here. **b)**. Run length. **c).** Run duration. The vast majority of the runs last for less than a second, but there are rare cases of these runs lasting tens of seconds. **d).** Stopping (tumbling) duration. As shown in the main text, these are exponentially distributed. Again, similar to the run duration, there are extremely rare cases where the tumbling duration will exceed tens of seconds.

**TABLE I.**
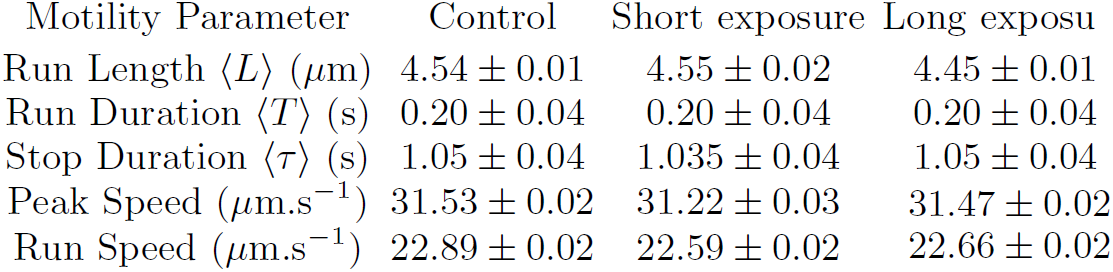
Mean values of the cell motility parameters for three different data sets with standard errors.

| Motility Parameter | Control | Short exposure | Long exposure |
| --- | --- | --- | --- |
| Run Length $\langle L \rangle$ ($\mu\text{m}$) | $4.54 \pm 0.01$ | $4.55 \pm 0.02$ | $4.45 \pm 0.01$ |
| Run Duration $\langle T \rangle$ (s) | $0.20 \pm 0.04$ | $0.20 \pm 0.04$ | $0.20 \pm 0.04$ |
| Stop Duration $\langle \tau \rangle$ (s) | $1.05 \pm 0.04$ | $1.035 \pm 0.04$ | $1.05 \pm 0.04$ |
| Peak Speed ($\mu\text{m.s}^{-1}$) | $31.53 \pm 0.02$ | $31.22 \pm 0.03$ | $31.47 \pm 0.02$ |
| Run Speed ($\mu\text{m.s}^{-1}$) | $22.89 \pm 0.02$ | $22.59 \pm 0.02$ | $22.66 \pm 0.02$ |

Similar to other marine organisms *Micromonas sp.* is an extremely fast swimmer relative to its body size - using high-speed microscopy techniques we have recorded rare cases where the peak speed of the cell exceeds 120 *µ*ms^−1^ or 60 body lengths per second. In the experiments conducted here the peak run speed is on average ≈ 32 *µ*ms^−1^ and is exponentially distributed as shown in Fig S3a. This large run velocity relative to the scale of the Brownian noise can be used to detect run events by examining the instantaneous speed of the particle as shown in Figure S2. From this information we can extract the x-y coordinates of the run trajectory, from which we can determine the value of the motility parameters for this run event.

Since we only consider straight runs, the run length distribution (Fig S3b) is the measured displacement of the cell during the run event. The run durations (Fig S3c) are exponentially distributed and typically last less than a second though we have observed rare events where the cell swims slower but for tens of seconds, able to cross the entire observation window. Finally, the tumble durations are also exponentially distributed (Fig S3d) as dscussed in the main text. Similar to the run durations, we observe rare cases of long tumble duration - with one event having a tumble time of over 200 s. For the average value shown in the main text, we only consider tumbles of less than 5 s since we can see from the distribution hat at this point the probability of a longer tumble is extremely low.

### SI 3: REORIENTATION ANGLE DISTRIBUTION

During the tumble phase of a run-tumble organism such as *Micromonas sp.*, the cell reorientates an angle *θ* measured counter clockwise from the previous run angle. During our cell motility experiments we are able to measure this rotation over a 2*π* period, but we are unable in these experiments to determine if the cell undergoes a number of complete rotations or not. Hence when fitting the probability distribution shown in the main text, we need to wrap a continuous probability distribution *P* (*θ*) over a unit circle, which produce the wrapped probability distribution as a 2*π* periodic sum.

Our derivation of the theoretical reorientation distribution goes as follows. Consider a run-tumble particle with an exponentially distributed tumble duration with characteristic time *τ*_*d*_:

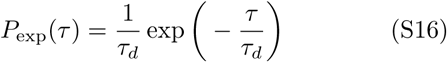

The particle is going to be subject to orientation from thermal forces with probability distribution function given as (with a peak reorientation angle of *θ*_*p*_ = 0):

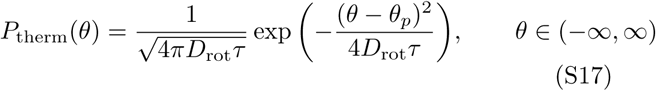

where *θ* is the reorientation of the particle during a time *τ* with rotational diffusion constant *D*_rot_. Combining these two distributions and summing over all values of the tumble duration gives the probability distribution:

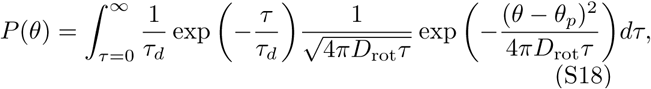

To apply this to *Micromonas sp.*, two modifications need to be made. Since we are only capable of experimentally measuring *θ* ∈ [−*π/*2, 3*π/*2], *P* (*θ*) needs to be wrapped to form the circular distribution *P*_*w*_(*θ*) by:

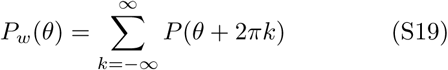

Secondly we need to account for the twin peaks of the experimental distribution of *θ*. This is done by combining two probability distribution with a parameter Γ, noting that if *P*_1_(*x*), *P*_2_(*x*) are normalized probability distributions then:

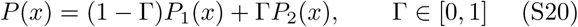

produces the normalized probability distribution *P* (*x*). Combining the two modifications of Eq S19, Eq S20 with our original probability distribution Eq S18, with *θ*_*p*_ = 0, *π*, we obtain:

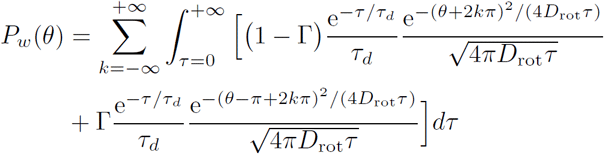

*P*_*w*_(*θ*) was fitted to the experimental data by minimising the Kullback-Leibler divergence, a measure of the divergence of two probability distributions. Given two probability distribution functions *p*(*x*), *q*(*x*) > ∀*x* ∈ *X*, it is possible to measure the divergence of the experimental data (*p*(*x*)) from the model distribution (*q*(*x*) = *P*_*w*_(*θ*)). As long as the above conditions are satisfied, the Kullback-Leibler divergence is defined as:

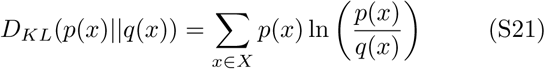

From this, we calculated the following parameter values for our fit: Γ = 0.55, *D*_rot_ = 2.1 rad.s^2^ and *τ*_*d*_ = 0.8 s. As we see in the main text, this distribution gives a very good description of the observed experimental data.

**FIG. S4.**
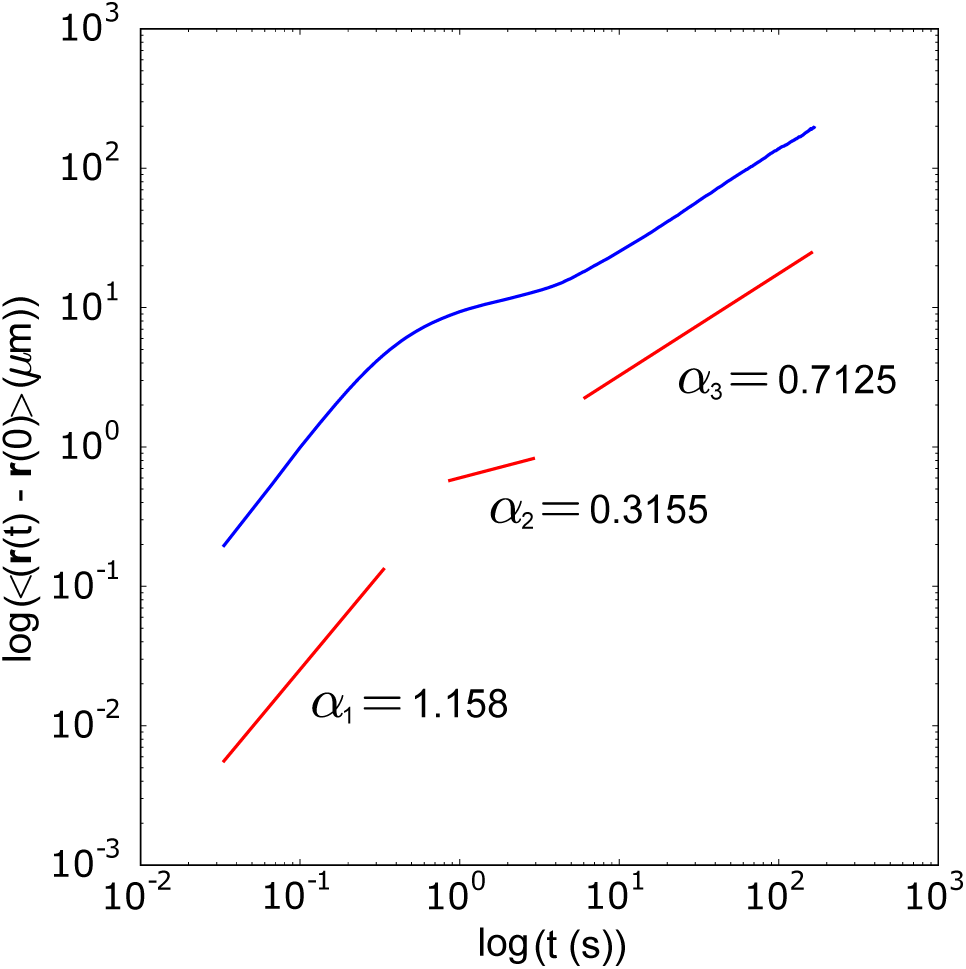
Average mean-squared displacement as a function of time. We observe three different diffusive regimes: an initial super-diffusive regime (*α*_1_ = 1.158), followed by a transition to a sub-diffusive regime (*α*_2_ = 0.3155, *α*_3_ = 0.7125) for long timescales.

### SI 4: DIFFUSION COEFFICIENT

The translational diffusion coefficient *D*_*t*_ was calculated from the averaged mean squared displacements (AMSD) of the particles, shown in Figure S4. The relationship between the AMSD and time for a two-dimensional system is defined as:

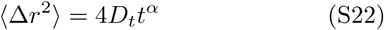

Here, we can see that setting *α* = 1 recovers the Brownian diffusion regime. In the case of anomalous diffusion, *α* > 1, *α <* 1 define the super-diffusive and sub-diffusive regimes respectively. We obtain three different regimes as shown in Figure S4; an initial super-diffusive regime followed by two sub-diffusive regimes with *α*_1,2,3_ = 1.158, 0.3155, 0.7125 respectively. While run-tumble styles of motion can lead to anomalous diffusion [S69, S70], we believe the subdiffusive behaviour observed for *Micromonas sp.* on long-time scales is in fact simply the consequence of our spatially-limited observation window (185*µ*m). We still extract the translational diffusivity via this long-time scale evolution of the AMSD leading to *D*_*t*_ = 1.17*µ*m^2^.s^−1^.

**FIG. S5.**
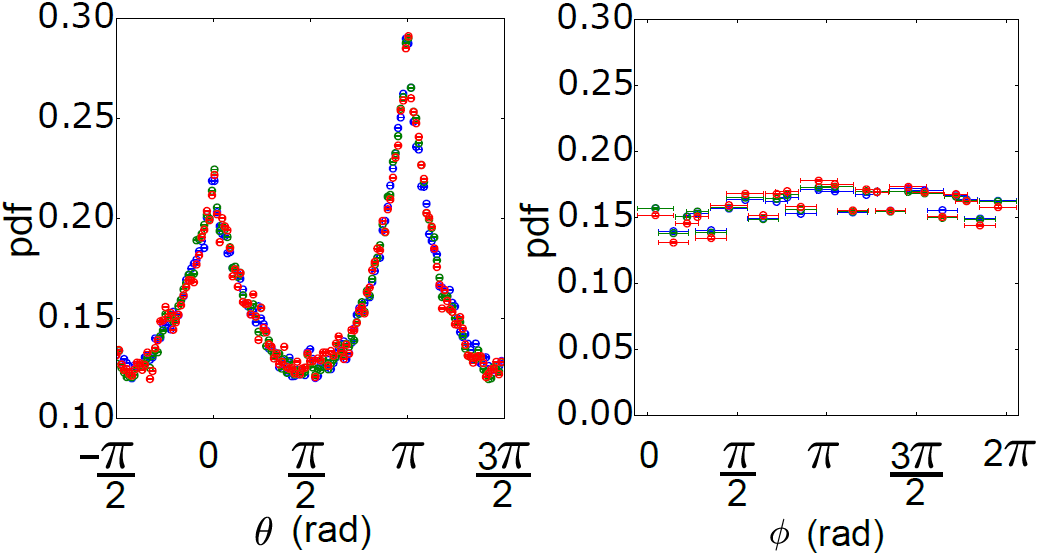
Angluar distributions with/without stimulus. In both figures, blue is the control, red the short-time stimulus and green the long-time stimulus. **a).** Reorientation-angle (*θ*). There is no change to the reorientation distribution when a stimulus is applied. **b).** Run-angle (*ϕ*) distribution. The distribution roughly uniform for both the control and stimulated cases, indicating no preferred run-direction.

### SI 5: RUN-ANGLE AND REORIENTATION-ANGLE DISTRIBUTIONS WITH/WITHOUT LIGHT STIMULUS

In previous run-and-tumble-taxis studies such as *E. coli* chemotaxis [S71], the chemotactic drift is produced by biasing the run-tumble durations to introduce a directional persistence to the motion. If this is the case here, it is expected that there will be a direct change to either the run-angle or reorientation-angle distributions (or both). For example, reducing tumbling duration should in theory increase directional persistence as the cell has less time to be reorientated by diffusion and so is more likely to travel towards/away the stimulus on the next run. For a kinesis-style of response, where the response is dictated by changing the run speed and/or run length, there would not necessarily be a change to either of these angular distributions. Figure S5 displays the probability distribution functions for the reorientation angle (*θ*, Fig S5a) and the run angle (*ϕ*, Fig S5b) in all three light conditions - control (no stimulus), short exposure and long exposure. We do not observe a significant difference in either distribution regardless of the presence of an external light stimulus.

### SI 6: SEDIMENTATION RATE

Dense samples of *Micromonas sp.* culture were loaded into a 50 *µ*m microfluidic chamber, inverted and left to sediment in a dirunal chamber. The samples were then flipped and the arrival times of the particles at the bottom surface recorded. Fig S6 plots the particle count *C*(*t*) (normalized by the maximum particle count attained, *C*_max_). After 52 min the count begins to plateau, resulting in a estimated sedimentation speed of 0.95 *µ*m.min^−1^, suggesting the slow phototactic drift is sufficient to maintain the cell’s vertical position in the water column and thus remain in optimal photosynthetic conditions.

**FIG. S6.**
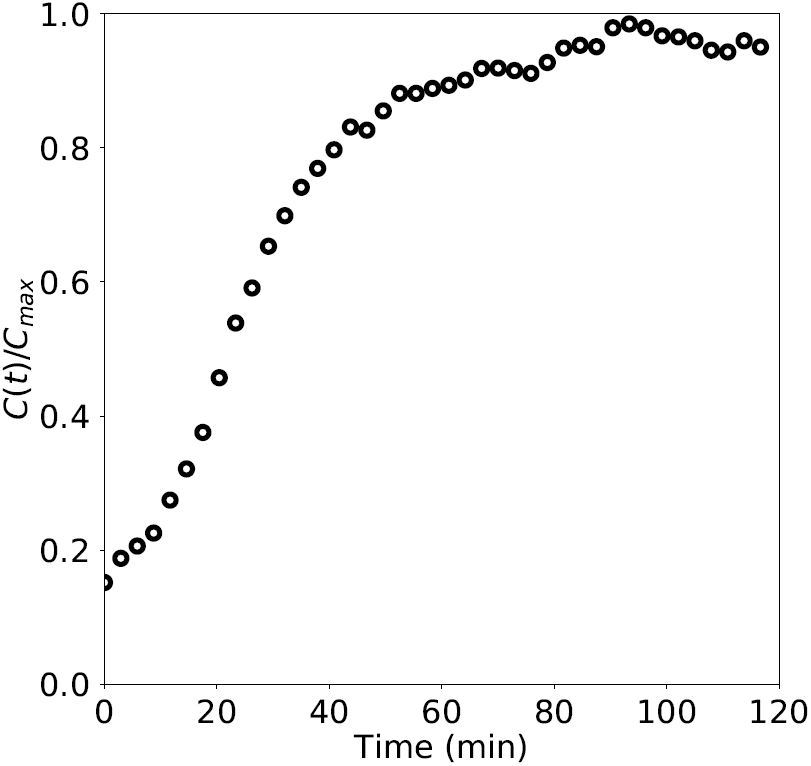
Sedimentation of *Micromonas sp.*. A dense sample of *Micromonas sp.* was loaded into a 50 *µ*m thick microfluidic chamber and inverted to collect particles on the top of the chamber. The chamber was then flipped and the arrival times of particles recorded, where *C*(*t*) is the number of particles in view at a time *t*, and *C*_max_ the maximum particle count attained. After 52.4 min the particle arrival begins to plateau, giving an estimate of 0.95 *µ*m.min^1^ for the sedimentation velocity.

### SI 7: CELL MICROLENSING

**FIG. S7.**
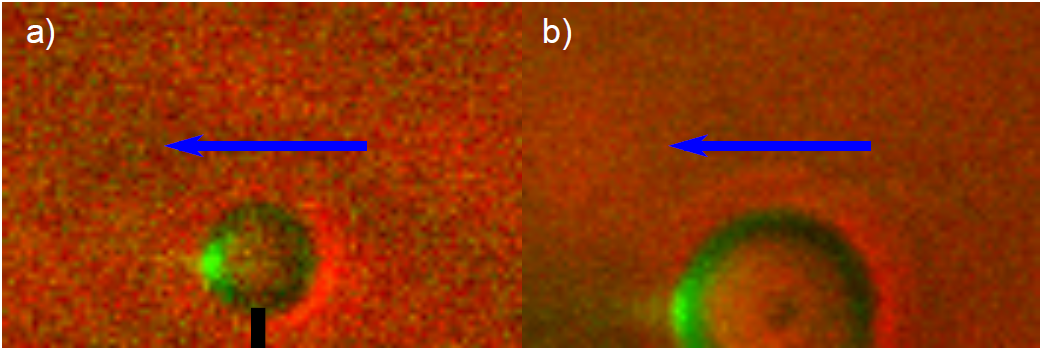
Microlensing in *Micromonas sp.*. Two different cells (a,b) in different orientations were illuminated with blue light directed parallel to the focal plane (direction of propagation: blue arrow) and imaged in both brightfield (red false colour) and through a red filter cube (green false colour). The latter captures the autofluorescence of the cells’ chloroplast, indicating strong focusing the external light.

*Micromonas sp.* cells were imaged with an 100× oil-immersion objective (with additional 1.5× internal magnification and illuminated with a blue (470 nm) LED perpendicular to the optical path and a FTIC cube was used to select the imaging wavelength, with the images recorded with an EMCCD camera. Fig S7 shows three different cells in different orientations (a-c) where the flagellum location is denoted by the black line, and the illumination direction by the blue arrow. In red is the brightfield image of the cell, with the fluorescent signal from the cell emission overlain in green. We see that the cell is capable of focusing external light internally in the cell suggesting a potential phototactic steering mechanism when the light is focussed onto light-sensitive components of the cell i.e. photoreceptors, chloroplast.

